# Frontal cortical regions associated with attention connect more strongly to central than peripheral V1

**DOI:** 10.1101/2020.09.09.281725

**Authors:** Sara Sims, Pinar Demirayak, Simone Cedotal, Kristina Visscher

**Affiliations:** University of Alabama Department of Psychology; University of Alabama Department of Neurobiology

**Keywords:** V1, structural connectivity, functional connectivity, visual eccentricity, functional networks

## Abstract

The functionality of central vision is different from peripheral vision. Central vision is used for fixation and has higher acuity that makes it useful for everyday activities such as reading and object identification. The central and peripheral representations in primary visual cortex (V1) also differ in how higher-order processing areas modulate their responses. For example, attention and expectation are top-down processes (i.e., high-order cognitive functions) that influence visual information processing during behavioral tasks. This top-down control is different for central vs. peripheral vision. Since functional networks can influence visual information processing in different ways, networks (such as the Fronto-Parietal (FPN), Default Mode (DMN), and Cingulo-Opercular (CON)) likely differ in how they connect to representations of the visual field across V1. Prior work indicated the central representing portion of V1 was more functionally connected to regions belonging to the FPN, and the far-peripheral representing portion of V1 was more functionally connected to regions belonging to the DMN.

Our goals were 1) Assess the reproducibility and generalizability of retinotopic effects on functional connections between V1 and functional networks. 2) Extend this work to understand structural connections of central vs. peripheral representations in V1. 3) Examine the overlapping eccentricity differences in functional and structural connections of V1.

We used resting-state BOLD fMRI and DWI to examine whether portions of V1 that represent different visual eccentricities differ in their functional and structural connectivity to functional networks. All data were acquired and minimally preprocessed by the Human Connectome Project. We identified central and far-peripheral representing regions from a retinotopic template. Functional connectivity was measured by correlated activity between V1 and functional networks, and structural connectivity was measured by probabilistic tractography and converted to track probability. In both modalities, differences between V1 eccentricity segment connections were compared by paired, two-tailed t-test. Dice Coefficients were used to determine spatial overlap between modalities.

We found 1) Centrally representing portions of V1 are more strongly functionally connected to frontal regions than are peripherally representing portions of V1, 2) Structural connections also show stronger connections between central V1 and frontal regions, 3) Patterns of structural and functional connections overlaps in the lateral frontal cortex.

In summary, the work’s main contribution is a greater understanding of higher-order functional networks’ connectivity to V1. There are stronger structural connections to central representations in V1, particularly for lateral frontal regions, implying that the functional relationship between central V1 and frontal regions is built upon direct, long-distance connections. Overlapping structural and functional connections reflect differences in V1 eccentricities, with central V1 preferentially connected to attention-associated regions. Understanding how V1 is functionally and structurally connected to higher-order brain areas contributes to our understanding of how the human brain processes visual information and forms a baseline for understanding any modifications in processing that might occur with training or experience.

## 1. INTRODUCTION

The function of central vision is different from peripheral vision. Central vision is used for fixation and has higher acuity, making it useful for reading and object identification (Larson & Loschky, 2009; Pelli et al., 2007; Trouilloud et al., 2020; Yoo & Chong, 2012). Peripheral vision has lower acuity but is important for visual tasks such as visual search and getting the gist of a scene (Larson & Loschky, 2009; Rosenholtz, 2016; Trouilloud et al., 2020). Differences in acuity between peripheral and central vision alone do not provide a full explanation of the extent of disparity in visual ability (Levi, Klein, & Aitsebaomo, 1985; Levi, Klein, & Wang, 1994). Information processing of central and peripheral visual information also differ. Peripheral visual information is processed faster than central information (Lu, Lesmes, & Dosher, 2002), and this fast processing helps determine objects’ salience within the visual field. This enables the visual system to direct saccades, fast eye movements, to a salient location (Lu et al., 2002). When central vision orients to the salient location, it provides high acuity information and distinguishes it from competing, distracting information (Lu et al., 2002).

The human primary visual cortex (V1) is organized along the calcarine sulcus in a progression of posterior representations of central vision (central V1) to anterior representations of peripheral vision (far-peripheral V1) (Duncan, Sample, Weinreb, Bowd, & Zangwill, 2007; Engel, Glover, & Wandell, 1997; Fox, Miezin, Allman, Van Essen, & Raichle, 1987). The anatomical regions representing central and peripheral V1 differ in their cortical thickness (Burge et al., 2016). The central representing portion of V1-- the part of the visual field used for fixation -- has a thicker cortex compared to the peripheral representing cortex in individuals with healthy vision (Burge et al., 2016).

There has been extensive prior research into the physiological differences between central and peripheral V1. Encoding of visual information differs within V1 representations of central and peripheral visual fields. The area of V1 devoted to central vision is much larger than that devoted to processing information from the peripheral visual field (Azzopardi & Cowey, 1993; Horton & Hoyt, 1991). In other words, the cortical magnification factor (square-mm of cortex devoted to each square-degree of visual angle) is greater for central vision than peripheral vision. Receptive field size is greater for neurons in peripheral V1 than in central V1 (Hubel & Wiesel, 1974). As eccentricity (distance from the center) increases, cortical magnification decreases, and receptive field size increases across human V1 and nearby visual areas (Harvey & Dumoulin, 2011).

Central and peripheral V1 also differ in how higher-order processing areas modulate their responses. Functional properties of cortical neurons are adaptive; top-down demands of high-order cognitive processing tasks influence their response (Gilbert & Li, 2013). For example, attention and expectation are top-down processes (i.e., high-order cognitive functions) that influence visual information processing during behavioral tasks (Gandhi, Heeger, & Boynton, 1999; Somers, Dale, Seiffert, & Tootell, 1999; Tootell et al., 1998; Yeshurun & Carrasco, 1998). This top-down control is different for central vs. peripheral vision. Attention influences the spatial summation of receptive fields so that spatial summation in foveal cells decreases and spatial summation in the peripheral cells in V1 increases (Roberts, Delicato, Herrero, Gieselmann, & Thiele, 2007). Psychophysiological data demonstrates that top-down influences on central vision stimuli are more potent than peripheral vision stimuli (Zhaoping, 2017). Similarly, attentional suppression of distractors is greater for central vision than peripheral vision (Chen & Treisman, 2008). Vision representations are known to interact strongly with higher-order brain networks (eg. (Casarsa De Azevedo, 2019; Furl, 2015; Gazzaley et al., 2007; Griffis, Elkhetali, Burge, Chen, & Visscher, 2015; Mantini, Corbetta, Perrucci, Romani, & Del Gratta, 2009; McMains & Kastner, 2011; Yeshurun & Carrasco, 1998), and these interactions appear to be distinct for central and peripheral vision.

Resting-state functional networks are groups of brain regions whose activity is temporally correlated at rest (Zalesky, Fornito, Cocchi, Gollo, & Breakspear, 2014). One way to identify a functional network is through clusters of correlated activity in the cortex (Yeo et al., 2011). The fronto-parietal network (FPN) has been shown to be important for directing attentional control (Zanto & Gazzaley, 2013), the cingulo-opercular network (CON) is involved in the maintenance of task demands (Coste & Kleinschmidt, 2016), and the default mode network (DMN) is less active when there are attentional or task goals (Raichle, 2015) and instead is thought to support tasks such as memory retrieval and semantic processing (Binder, 2012; Gerlach, Spreng, Gilmore, & Schacter, 2011; Sestieri, Shulman, & Corbetta, 2010; Spreng, 2012). Since functional networks can influence visual information processing in different ways, networks likely differ in how they connect to representations of the visual field across V1. There are also feedforward connections between V1 and functional network regions, as evidenced by the fact that strong stimulus-driven signals are observed in higher-order brain regions (Katsuki & Constantinidis, 2014).

Previous work in our lab has investigated how the central-to-peripheral cortical organization of V1 influences functional connectivity between V1 and the rest of the cortex (Griffis et al., 2017). This prior work showed retinotopic patterns of functional connectivity between V1 and functional networks during resting fixation. Specifically, the central representing portion of V1 was more functionally connected to regions belonging to the FPN, and the far-peripheral representing portion of V1 was more functionally connected to regions belonging to the DMN. The fact that, as described above, central vision is under different top-down control than peripheral vision might underlie its preferential connection to the FPN, which is related to directing attentional control and has been shown to facilitate bottom-up and top-down attentional processes for visual information (Katsuki & Constantinidis, 2014). The role of far-peripheral vision in environmental monitoring and the need to suppress visual information from this portion of the visual field during central fixation might be the reason for preferential connectivity to the task-negative DMN (Li, 2002). However, this previous work was limited by a small sample size (i.e., 20 participants) and design (i.e., data were acquired during the resting fixation blocks of a visual task) issues (Griffis et al., 2017). These limitations leave questions regarding whether previous functional findings extend to free viewing during rest in a larger sample and, critically, how these effects relate to differences in the anatomical (i.e., white matter) connections of central vs. peripheral V1.

Anatomical connections can be examined by an invasive method in which retrograde or anterograde tracers are injected into the brain, and the resulting staining of brain regions is examined. These studies inject one location and look for staining in other locations, giving finely detailed information about the brain’s connection patterns (Felleman & Van Essen, 1991). Because of the slow and arduous nature of these studies, examinations of visual regions’ connectivity have primarily focused on defining structural connectivity across areas thought to be primarily involved with vision (Andersen, Asanuma, Essick, & Siegel, 1990; Lysakowski, Standage, & Benevento, 1988; Neal, Pearson, & Powell, 1990). Thus, it is unclear whether prior work did not find connections between V1 and higher-order processing areas or simply did not have the capability to observe those findings from their experiments. A more comprehensive study aimed to close this literature gap by creating a connectivity matrix throughout the macaque’s cerebral cortex (Markov et al., 2014). Although we cannot make explicit homologies between human resting state networks and cortical areas in macaque, we can say that a limited number of frontal and parietal areas were examined, and these may be homologs to components of the FPN. An injection of anterograde tracer placed into central-representing V1 resulted in the identification of connections projecting from V1 to regions of the frontal and parietal cortex including F5 (a frontal area involved in motor planning), 8l (frontal eye fields), and 7A (a parietal area involved in attention modulation and planning) (Markov et al., 2014). Of the frontal and parietal areas examined by Markov et al., only area 8l (frontal eye fields) showed top-down connections to V1. Significantly, in the taxonomy put forward by Yeo and colleagues (2011), the “fronto-parietal network” does not overlap with frontal eye fields (Glasser, Coalson, et al., 2016; Majerus, Péters, Bouffier, Cowan, & Phillips, 2018; Wang, Mruczek, Arcaro, & Kastner, 2015). Thus there were no top-down connections to V1 from candidate fronto-parietal regions in the Markov paper. There were no tracers injected into the peripheral representing portion of V1. Thus, it is not clear whether these candidate bottom-up V1-fronto-parietal connections (areas F5 and 81) exist for peripheral portions of V1.

While brain anatomy is relatively fixed in adulthood, a healthy brain adapts with changes in structural connections between regions with experience and age (Davis et al., 2009). White matter tracts correspond to direct connections between brain regions. Structural connections can be studied in-vivo in humans with diffusion-weighted MRI and tractography. Major white matter tracts that connect to the occipital lobe, such as the inferior fronto-occipital fasciculus (connects occipital lobe to the lateral prefrontal cortex) and the inferior longitudinal fasciculus (connects occipital lobe to anterior temporal lobe), have been well documented using tractography methods in humans (Wu, Sun, Wang, & Wang, 2016). However, the inferior fronto-occipital fasciculus has not been found in the macaque brain, which may explain why tracer studies investigating V1 have not helped inform occipital-prefrontal cortex connections (Takemura et al., 2016).

The current study’s goals are 1) to assess the reproducibility and generalizability of retinotopic effects on functional connections between V1 and functional networks found in prior work (Griffis et al., 2017). We aim to extend these findings in a new dataset collected under different task conditions (previous work used blocks of rest during a task with central fixation, and the current data was collected as part of a resting-state only scan). 2) Extend prior work on the retinotopic connectivity difference to structural connections between V1 and functional networks. 3) Examine regions of overlap between functional and structural connections. Since functional connectivity between two brain regions could reflect measurable structural connections, we used DWI to examine connections between regions (Adachi et al., 2012; Honey et al., 2009).

To address these goals, we used resting-state BOLD fMRI and DWI to examine if portions of V1 that represent different visual eccentricities differ in their functional and structural connectivity to functional networks. We found 1) substantial evidence that centrally representing portions of V1 are more strongly functionally connected to lateral frontal regions than are peripherally representing portions of V1, 2) Structural connections show the same pattern, with stronger connections between central V1 and frontal regions, in particular a lateral frontal portion of the FPN, and 3) the pattern of structural and functional connections is similar, suggesting that this lateral frontal functional connection pattern arises from a direct (uni-synaptic) structural connection. These results, coupled with relationships to prior work described in the discussion, are suggestive that the processing of central vision is mediated in part through direct connections to the lateral frontal cortex.

## 2. METHODS

### 2.1 Participants

The study used diffusion-weighted imaging, resting-state functional imaging, and structural imaging data from the 900-subject release of the Human Connectome Project (HCP) dataset (Figure 1). Participants in this dataset were healthy young adults between 22-36 years of age who had normal or corrected-to-normal vision. Most subjects had at least one relative in the group; many of them are twins. Our hypotheses are not about individual differences, and due to the large sample size of the data, there is still a great deal of diversity in the sample; therefore, we did not treat related and unrelated samples separately. Using this relatively large sample size facilitates replication and extension of findings.

**Figure 1.**
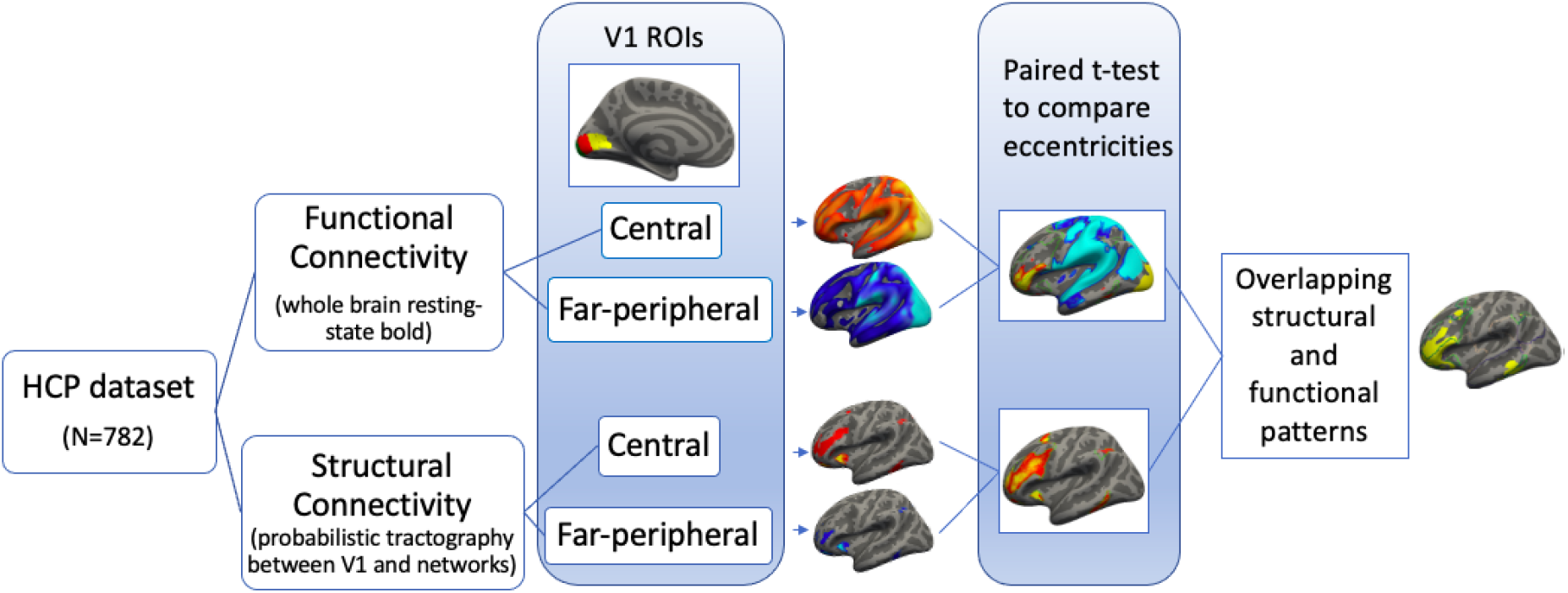
Graphical representation of methods. Moving from left to right: data from the HCP dataset through the analysis stream and representations of the results. The top portion shows the analyses for functional connectivity, and the bottom portion shows the analysis for structural connectivity. The overlap between the two modalities was calculated with a dice coefficient.

We excluded participants with structural abnormalities (e.g., tumors and extensive area brain damage) identified through HCP quality control. We then visually inspected the remaining data for white matter abnormalities. We excluded participants if their structural scans displayed large or punctate white matter hyperintensities easily detected by eye. In total, we excluded 114 subjects from the original 900 subject dataset. Seven hundred eighty-six healthy subjects passed quality control standards, including 335 males and 449 females (Figure 1; see participant IDs in code repository). We excluded the fMRI data from four of them due to quality standards for functional comparison. Otherwise, we used the same participants in both structural and functional connectivity analyses.

### 2.2 Data Acquired by the Human Connectome Project

All data were acquired and minimally preprocessed by the Human Connectome Project (Glasser, Smith, et al., 2016; Sotiropoulos et al., 2013; David C Van Essen et al., 2013; D C Van Essen et al., 2012). Informed consent was obtained by the Human Connectome Project (David C Van Essen et al., 2013). The IRB for the University of Alabama at Birmingham reviewed our use of the HCP dataset and determined it exempt.

T1-weighted structural MRI, resting state fMRI and multi-shell diffusion weighted (DWI) MRI data were acquired using a customized Siemens 3T “Connectome Skyra” (Sotiropoulos et al., 2013). High-resolution three-dimensional MPRAGE, T1-weighted anatomical images (TR = 2400 ms, TE = 2.14 ms, flip angle = 8, FOV = 320 x 320 mm^2^, voxel size 0.7 x 0.7 x 0.7 mm^3^, number of slices = 256, acceleration factor (GRAPPA) = 2) were used. Functional magnetic resonance imaging (fMRI) data were acquired with a multi-band gradient-echo (GE) EPI sequence (voxel size = 2 x 2 x 2 mm^3^; TR= 720 ms; TE = 33.1 ms; flip angle = 52 degrees; FOV = 208 x 180 mm^2^; number of slices = 72) in four runs (each of them took approximately 15 minutes) with eyes open and related fixation on a cross on a dark background. Phase encoding direction was right-to-left for half of the scans and left-to-right for the other half of the resting-state scans.

For DWI, multi-band diffusion-weighted echo-planar (EP) images (voxel size = 1.25 x 1.25 x 1.25 mm^3^; TR= 5520 ms; TE = 89.5 ms; flip angle = 78 degrees; MB = 3; FOV = 210 x 180 mm^2^; number of slices = 111; b=1000, 2000 and 3000 s/mm^2^, diffusion directions = 95, 96 and 97) were used. DWI data includes six runs (each of them took approximately 9 minutes and 50 seconds). Each gradient table was acquired with right-to-left and left-to-right phase encoding polarities which were then merged after distortion correction as part of the HCP Preprocessing Pipeline (Glasser et al., 2013).

### 2.3 V1 Eccentricity Segment Definitions

V1 eccentricity segments were hand-drawn within the Freesurfer fsaverage V1 label as described in a previous publication from our lab (Burge et al., 2016; Griffis et al., 2017, 2015). Previous work has shown that cortical anatomy is a reliable predictor of the retinotopic organization of V1 (Benson et al., 2012; O. Hinds et al., 2009; O. P. Hinds et al., 2008) so that the more posterior parts of the visual cortex represent more central portions of the visual field. The average eccentricity of each segment was estimated from Benson and colleagues’ retinotopy template (Benson, Butt, Brainard, & Aguirre, 2014; Benson & Winawer, 2018). Based on this template, we identified three retinotopic regions: central vision (mean eccentricity estimates of 0-2.2 degrees visual angle), mid-peripheral vision (mean eccentricity estimates of 4.1-7.3 degrees visual angle), and far-peripheral vision (mean eccentricity estimates of 14.1-25.5 degrees visual angle) (Figure 2).

**Figure 2.**
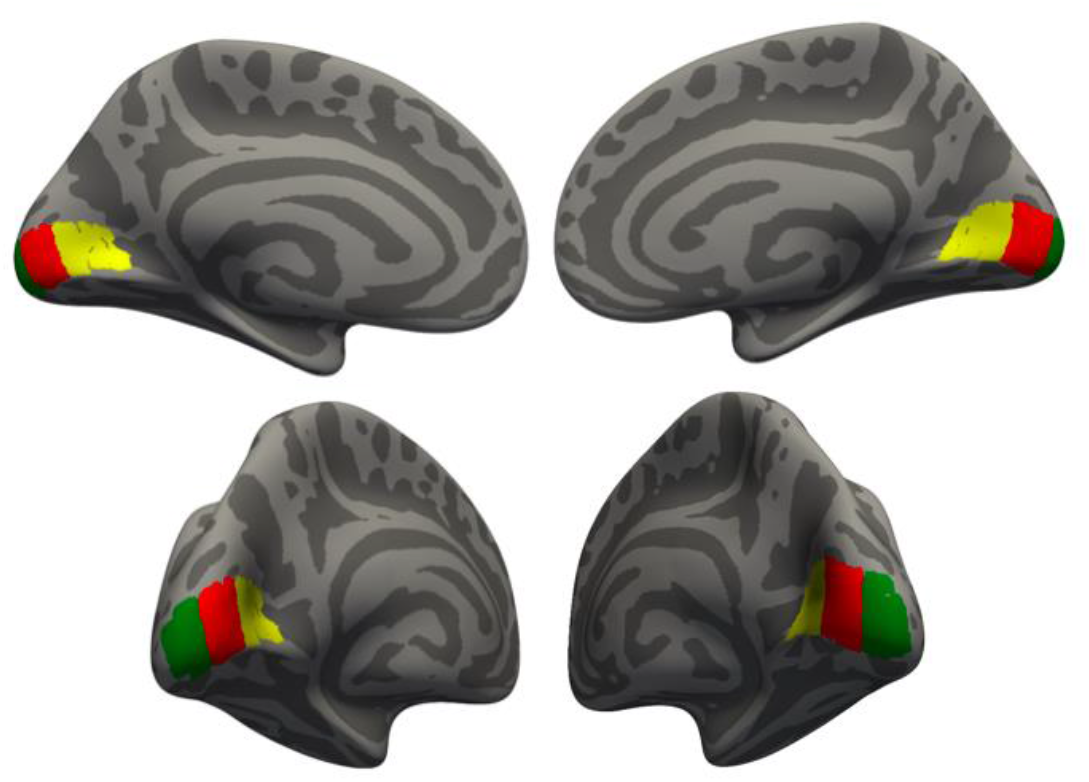
V1 Eccentricity segments. The Far-peripheral representing section of V1 is shown in yellow, the mid-peripheral representing section of V1 is shown in red, and the central representing section of V1 in green. These regions of V1 were used as seed regions in functional connectivity analyses. Central and far-peripheral ROIs were used as target regions for tractography analyses. For brevity, we sometimes use shorthands like “central V1” to refer to these centrally-representing regions.

These V1 eccentricity segment ROIs were defined on FreeSurfer’s fsaverage brain using the retinotopic template’s eccentricity, and then we interpolated them to the individual subjects’ cortical surfaces using FreeSurfer’s anatomical registration.

### 2.4 Functional Network ROI Definitions

We transformed the FPN, CON, and DMN labels created by (Yeo et al., 2011) from the Freesurfer fsaverage brain to individual anatomical space (Yeo et al., 2011). We used voxels within the grey matter corresponding to the network-ROIs as seed voxels for the functional connectivity analysis. We used voxels within the white matter corresponding to the network-ROIs as track seeds for the probabilistic tractography analysis. Voxels were identified using the Freesurfer mri_aparc2aseg command and then transformed into individual diffusion space.

### 2.5 Data Analysis

#### 2.5.1 Resting-State Scan Image Preprocessing

The HCP minimal preprocessing pipeline that includes artifact removal, motion correction, and registration to common space was used (Fischl, 2012; Glasser et al., 2013; Mark Jenkinson, Bannister, Brady, & Smith, 2002; M Jenkinson, Beckmann, Behrens, Woolrich, & Smith, 2012; D C Van Essen et al., 2012). Along with the preprocessing steps already described by Glasser and colleagues (2013), we applied additional preprocessing steps on the residual BOLD data to reduce spurious variance not associated with the neural activity as described in this paragraph. We then censored the functional images for movement according to validated techniques (Carp, 2013; Griffis et al., 2017; Power, Barnes, Snyder, Schlaggar, & Petersen, 2012). We replaced time points in which a participant moved more than 0.5 mm in one TR with an interpolated image from adjacent images. We excluded runs if the mean framewise displacement across the run was greater than 3 mm in any direction. We applied temporal band-pass filtering between 0.009 and 0.08 Hz. We applied regressors to reduce artifactual noise, including white matter and CSF signals and motion parameters that we extracted during motion correction for each subject from the previous step. Surface reconstruction, the region of interest (ROI) label generation, and image registration were also visually inspected for all subjects to ensure the automated computations’ accuracy. Next, we concatenated both the acquisitions (those collected right-to-left and those collected left-to-right) into a single 4D volume for the functional connectivity analysis.

#### 2.5.2 Functional Connectivity Analysis

Functional connectivity refers to synchronization between time courses of activation between two brain areas due to the similar temporal signal profiles from these connected areas (Friston, Holmes, Poline, Frith, & Frackowiak, 1995). Correlation maps for each participant were obtained from seed-to-voxel connectivity measurements between central, mid-peripheral, and far-peripheral ROIs within the primary visual cortex (V1) to each voxel in the brain (Figure 3). The resulting correlation coefficient maps were converted to z-score maps using Fisher’s z transform. Fischer’s transformed z-score maps were projected onto the individual cortical surface from 1mm below the white/gray matter boundary using Freesurfer’s mri_vol2surf command. We compared difference maps by paired, two-tailed t-test using Freesurfer’s mri_glmfit function to test the functional connectivity differences.

**Figure 3.**
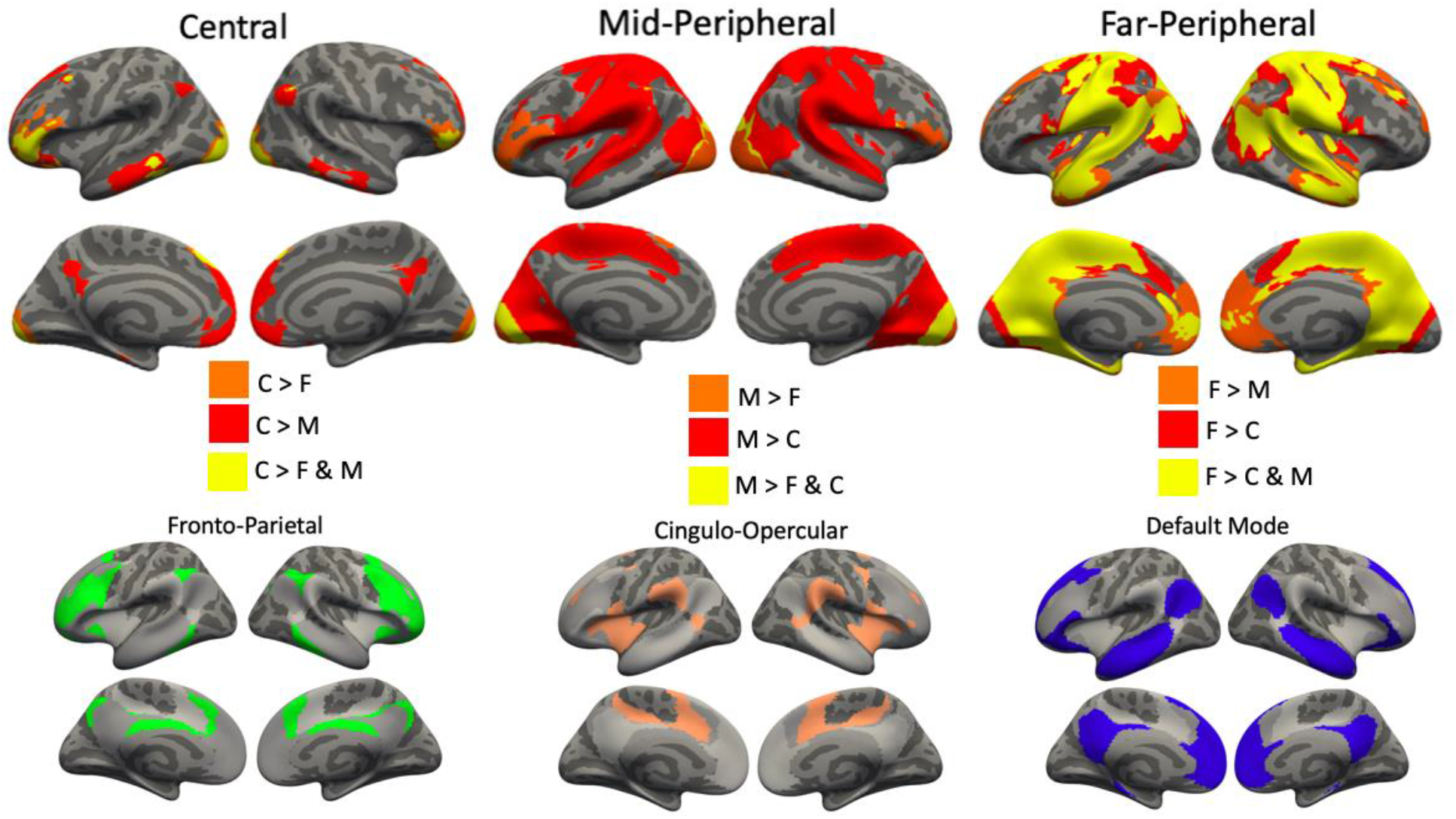
Comparisons of functional connectivity between V1 eccentricity segments and homology to known resting-state networks. *Top Row*: Differences in Functional Connectivity, depending on eccentricity. The far left panel highlights vertices with significantly stronger connections to central V1 than other portions of V1. The yellow vertices showed stronger connectivity to central V1 than to both far-peripheral and mid-peripheral regions. Red indicates stronger connectivity to central than mid-peripheral regions, and orange indicates stronger connectivity to central than far-peripheral regions. The middle and right panels show similar images highlighting vertices with significantly stronger connections to mid-peripheral and far-peripheral regions. For the middle panel, red is where mid-peripheral is greater than central, and orange is where mid-peripheral is greater than far-peripheral. For the panel on the right, red is where far-peripheral is greater than central, and orange is where far-peripheral is greater than mid-peripheral. We based Inferences regarding functional connectivity on these maps. *Bottom Row*: Functional Networks for comparison to the top-row. Previously documented FPN, CON, and DMN (from Yeo et al., 2011). The FPN is shown in green, the CON is shown in tan, and the DMN is shown in blue. The grey regions indicate the location of the other networks. Note homologies between the FPN and the top row left panel.

#### 2.5.3 Diffusion-weighted Image Preprocessing

The HCP minimal preprocessing pipeline was used to correct B0 and eddy current distortions (Andersson, Skare, & Ashburner, 2003; Andersson & Sotiropoulos, 2015, 2016; Glasser et al., 2013; Glasser, Smith, et al., 2016; Sotiropoulos et al., 2013). Further, we performed DWI data preprocessing using the FMRIB’s Diffusion Toolbox (FDT v3.0) using GPU for the acceleration of processing (graphics processing unit) (Hernández et al., 2013; Robinson et al., 2018). We estimated the distribution of diffusion parameters using Markov Chain Monte Carlo sampling for each voxel, allowing for crossing fiber orientations (Behrens, Johansen-Berg, Jbabdi, Rushworth, & Woolrich, 2007).

#### 2.5.4 Tractography

The results from our previous study on functional connectivity, replicated here, led to the hypothesis that structural connections from central vs. peripheral regions would differ between the FPN, CON, and DMN. Thus, we used these network regions as seeds in probabilistic tractography performed by FMRIB’s Diffusion Toolbox (FDT) (Hernandez-Fernandez et al., 2019) and used the V1 ROIs as targets. For each seed voxel, we calculated 10,000 streamlines (along with default settings of maximum steps: 2000, step length: 0.5mm, curvature threshold: 0.2) and separate samples of the voxelwise diffusion distribution. A distance correction and loop-check, which prevents circular pathways, were applied. The tractography then resulted in each voxel within the seed ROI containing the number of streamlines that reached the target (V1 region) from that voxel. We performed tracking in individual diffusion space. We transformed seed and target regions into diffusion space for tractography analysis, and then we transformed tractography results into individual anatomical Freesurfer space for visualization. Surface maps of the track termination probabilities were smoothed using a 2mm FWHM Gaussian filter and averaged across all subjects.

We transformed track frequencies (number of streamlines that reached the target) into track probabilities (likelihood of a track reaching the target) by dividing the log-scaled track frequency by the maximum log-scaled track frequency (Beer, Plank, & Greenlee, 2011; Wirth, Frank, Greenlee, & Beer, 2018). Track probabilities mitigate possible biases arising from size differences of seeds (Smith, Beer, Furlan, & Mars, 2018; Wirth et al., 2018). Track probabilities were projected onto the individual cortical surface from 1mm below the white/gray matter boundary using Freesurfer’s mri_vol2surf command (Beer et al., 2011; Wirth et al., 2018). Surface maps of the track termination probabilities were smoothed using a 2mm FWHM Gaussian filter and averaged across all subjects.

#### 2.5.5 Tractography Analysis

To statistically test patterns of structural connections, we compared the central and far-peripheral eccentricity segments of V1 connectivity patterns within the FPN, CON, and DMN. Differences in track probabilities corresponding to V1 eccentricity segment connections were compared by paired, two-tailed t-test (using Freesurfer’s mri_glmfit with a one-sample group mean test) (Figure 5).

#### 2.5.6 Comparison of Functional and Structural Connectivity

A Dice Coefficient calculates the spatial overlap between two spatial maps (Novosad, Fonov, Collins, & Alzheimer’s Disease Neuroimaging Initiative†, 2020; Zou et al., 2004). To compare structural and functional connectivity patterns of central vs. far-peripheral V1, we used the Dice coefficient to compare structural and functional maps within the FPN, CON, DMN, and across all 3 of the networks together. The functional map used in these analyses were the functional connectivity differences between central and far-peripheral V1 ROIs, thresholded at p<0.001 (See Figure 4). The structural map used in these analyses were the structural connectivity differences between central and far-peripheral V1 ROIs thresholded at p <0.001 (See Figure 6). We used a p-value threshold of p<.001 to include only those vertices with confidence in the differences’ direction. Data from the two hemispheres were calculated separately and then averaged together.

**Figure 4.**
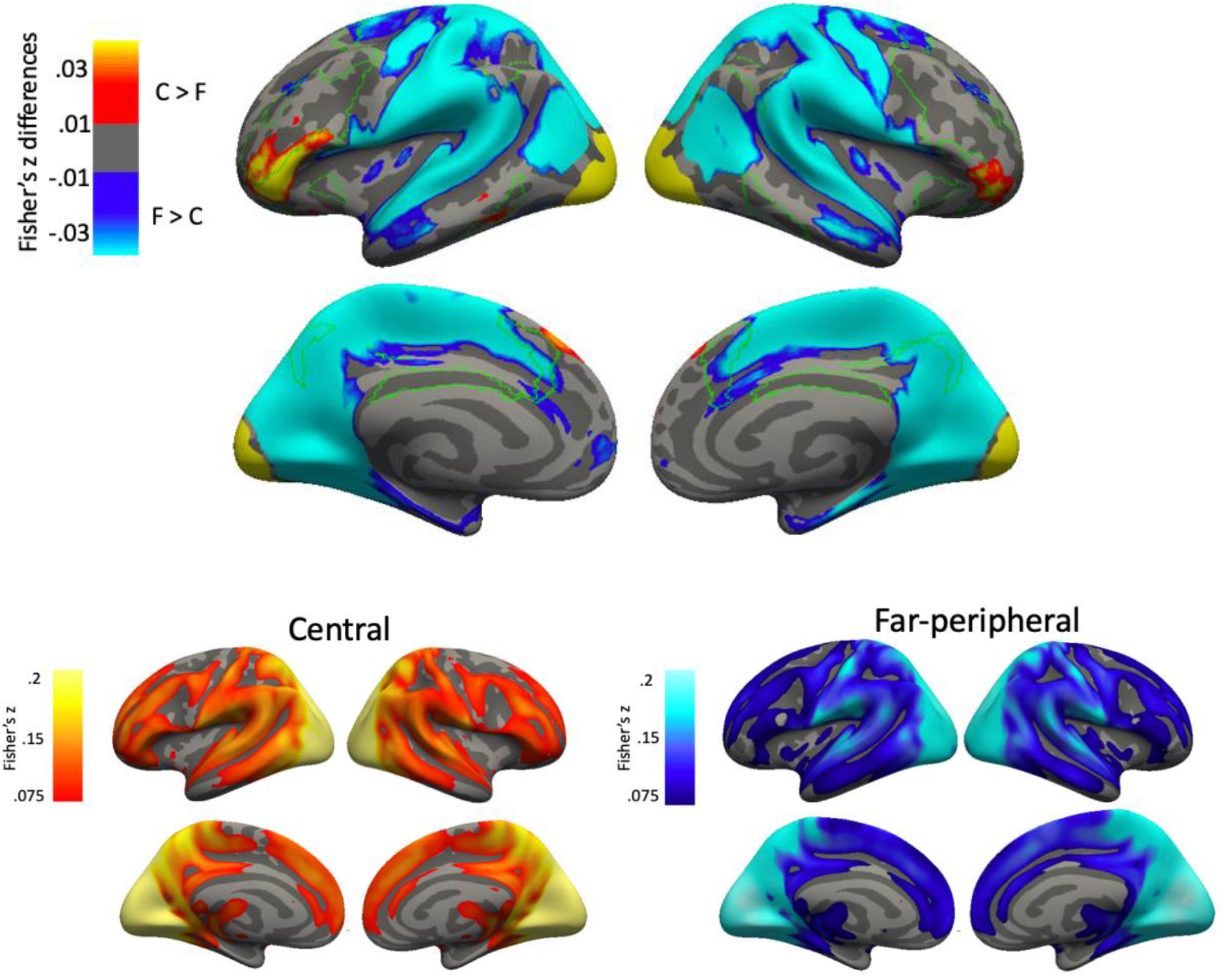
Group average and statistical maps of comparisons of functional connectivity between V1 central and far-peripheral segments. *Top: Group average central minus far-peripheral differences in functional connections with the FPN*. Group average data was thresholded for significance (p<.001) and effect size (connectivity differences > .01). The FPN is outlined in green.*Bottom: Group average for central and peripheral functional connectivity. The top image is a subtraction between the data from the bottom two images*.

**Figure 5.**
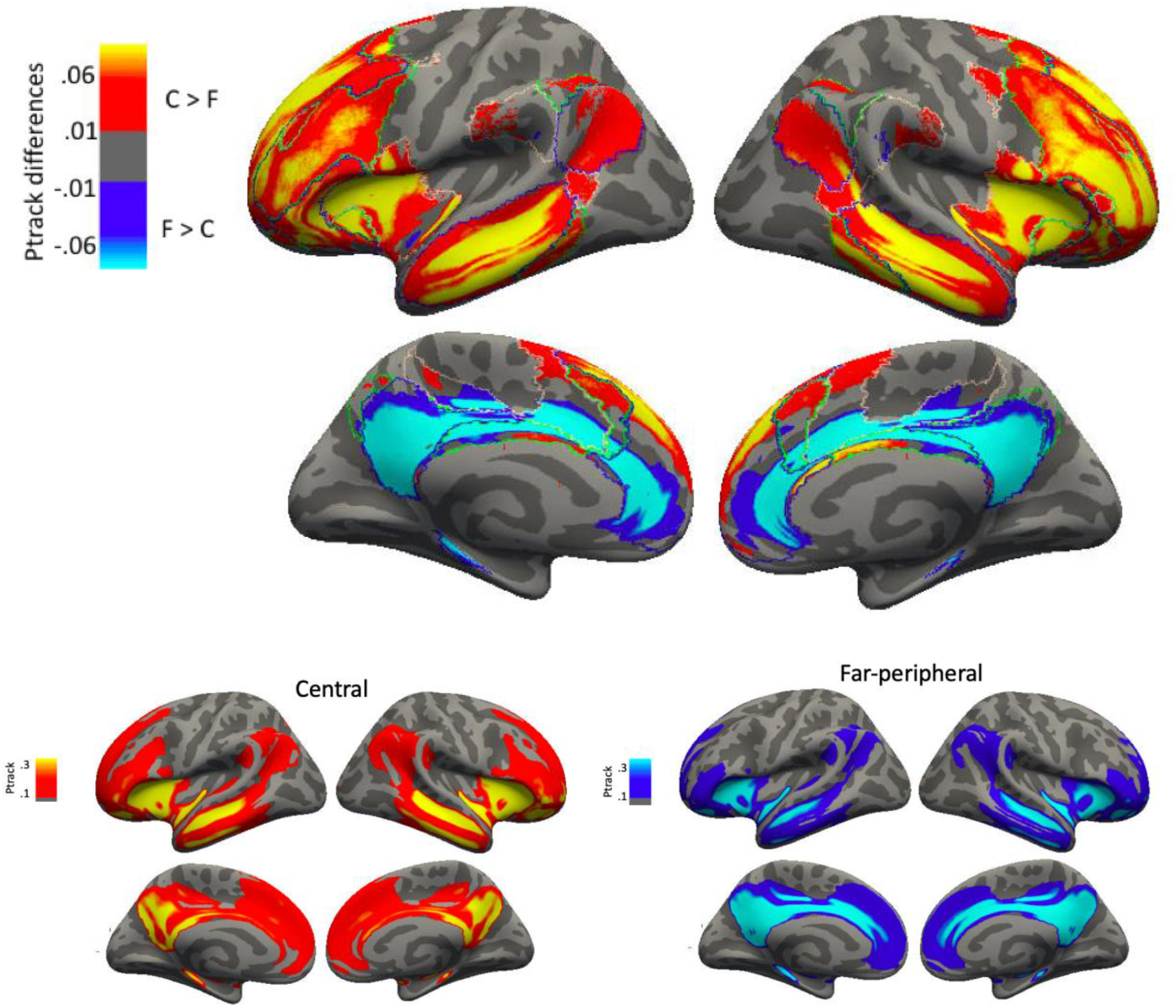
Group average and statistical maps of comparisons of structural connectivity between V1 eccentricity central and far-peripheral segments. *Top: Central (hot) minus Far-peripheral (cool) V1 structural connection differences to vertices within the FPN, CON, and DMN*. Group average p-track differences between central and far-peripheral V1 data were thresholded for significance (p<.001) and effect size (ptrack differences > .01) and masked for the FPN (outlined in green), DMN (outlined in dark blue), and CON (outlined in beige). *Bottom: Group average for central and peripheral structural connectivity. The top image is a subtraction between the data from the bottom two images*.

**Figure 6.**
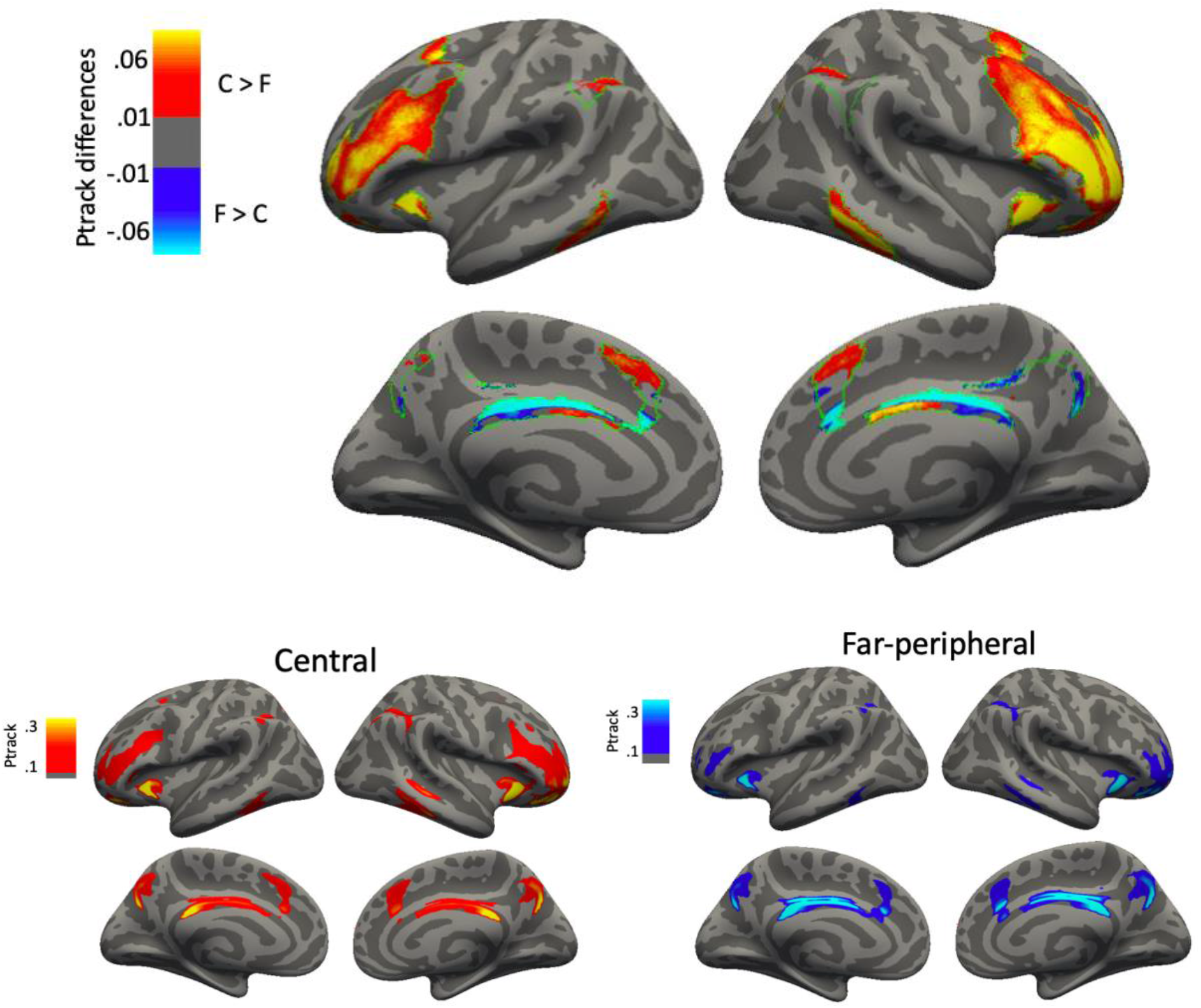
Group average and statistical maps of comparisons of structural connectivity between V1 eccentricity central and far-peripheral segments. *Top: Central minus Far-peripheral V1 structural connection differences within the FPN*. Group average p-track differences between central and far-peripheral V1 data were thresholded for significance (p<.001) and effect size (ptrack differences > .01) and masked for the FPN (outlined in green). *Bottom: Group averages for central and peripheral connectivity*.

#### 2.5.7 Correspondence between Functional Networks and Multi-modal Connectivity Patterns

We defined a multi-modal connectivity pattern as the voxels where connectivity for central V1 was greater than far-peripheral V1 (t-test, threshold p < 0.001) in *both* the structural and functional data (Figure 7). We calculated the Dice coefficient comparing this multi-modal connectivity pattern to each of the functional network ROIs (FPN, CON, DMN). These Dice coefficients quantify the overlap between the mulit-modal connectivity pattern and these network regions.

**Figure 7.**
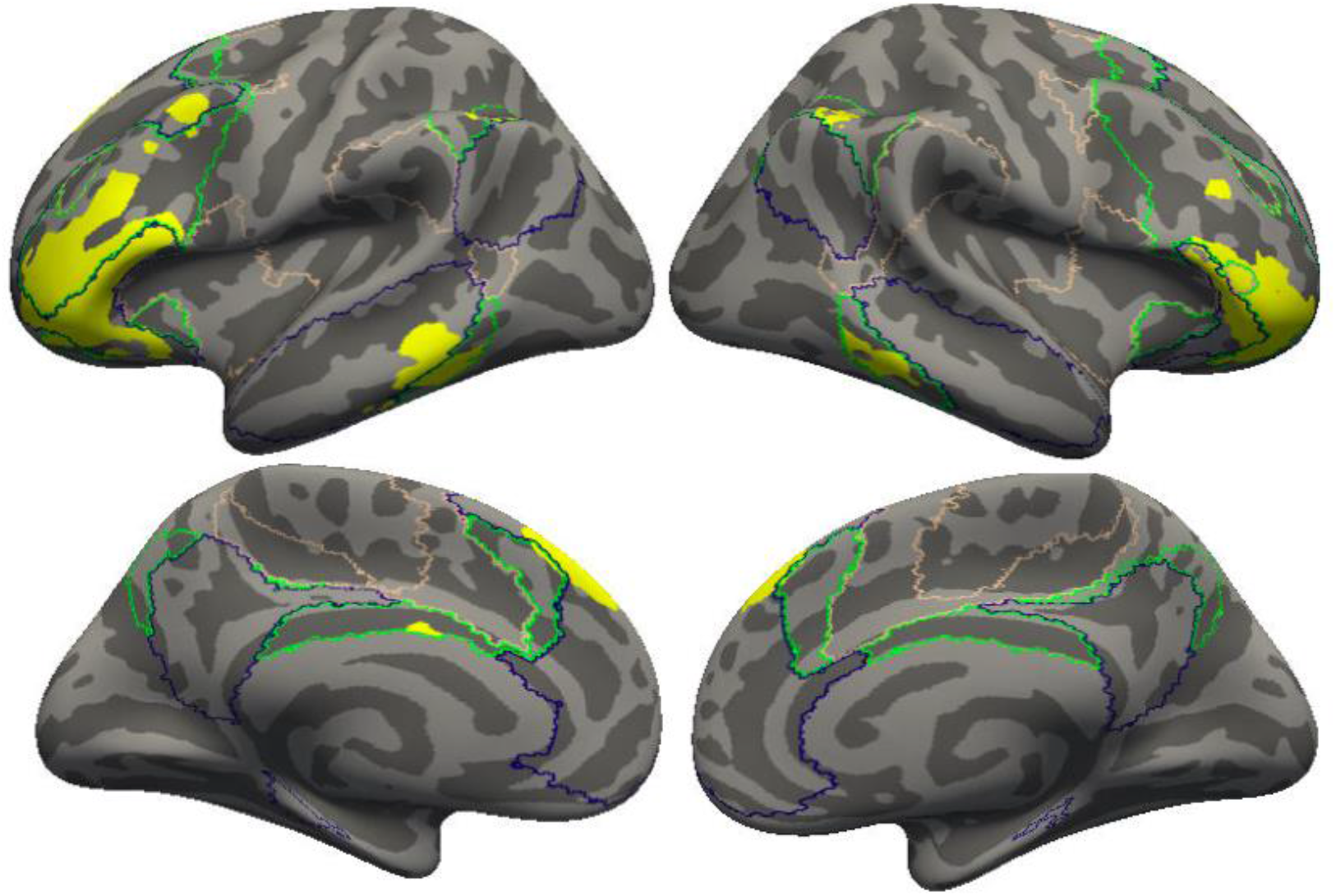
**Overlap of functional and structural connectivity patterns of central greater than far V1 eccentricity segments within resting-state networks (FPN (green outline), CON (beige outline), and DMN (dark blue outline))**.

## 3. RESULTS

We hypothesized that the connectivity between the eccentricity segments of the primary visual cortex (V1) and functional networks (i.e., FPN, CON, DMN) differs in both structural and functional connections.

### 3.1 Functional Connections to V1 depend on Eccentricity

We compared the whole-brain functional connectivity patterns of each segment of V1-central, mid-peripheral, and far-peripheral. The t-test comparing functional connectivity to different eccentricity segments in V1 revealed significant effects (*p*<.001) in brain regions belonging to FPN, CON, and DMN functional networks (Figure 3). Notably, central representing V1 was preferentially connected (over mid-peripheral and far-peripheral V1) to regions associated with the FPN, including the mid orbitofrontal and inferior parietal regions of the FPN (Figure 3, left). Mid and far-peripheral representing V1 were not preferentially connected (over central V1) to specific networks (Baldassano, Fei-Fei, & Beck, 2016). This finding of patterns of voxels with preferential connectivity to central V1 is similar to those found in a previous publication from our lab (Griffis et al., 2017). However, the patterns of voxels with preferential connectivity to mid-peripheral and far-peripheral V1 were not as distinct from each other as in the other dataset (Figure 3 middle and right). Thus, in this paper, we focus on distinctions between centrally-representing portions of V1 to far-peripheral portions of V1.

Due to indications of similarity between patterns of functional connectivity of central V1 and the FPN (from previous data in Griffis et al., 2017), we directly compared central and peripheral V1 functional connections to FPN. We performed pairwise statistical comparisons (t-test) of functional connections between central vs. far-peripheral eccentricity segments of V1 and the FPN (Figure 4). Results indicate that, like our initial functional connectivity findings (Figure 3), there are preferential connections between central V1 and the inferior frontal gyrus compared to far-peripheral V1 (Figure 4). This inferior frontal gyrus region aligns well with the anterior portion of the FPN as defined by Yeo, but interestingly, it does expand somewhat beyond that border into more inferior parts of the Inferior Frontal Gyrus (IFG). Like the FPN, the IFG has been shown to be related to attention and control (Baldauf & Desimone, 2014; Chong, Williams, Cunnington, & Mattingley, 2008; Fassbender et al., 2004; Hampshire, Chamberlain, Monti, Duncan, & Owen, 2010; Swick, Ashley, & Turken, 2008, 2011).

### 3.2 Structural Connectivity Eccentricity Differences

Next, we investigated similar comparisons between central and far-peripheral V1 in a different modality-structural connections. A t-test comparing the structural connections of central and far-peripheral V1 revealed significant effects (p<.001) in brain regions belonging to FPN, CON, and DMN functional networks (Figure 5). We chose these three networks to compare to functional connectivity findings from Figure 3.

Notably, central representing V1 was preferentially connected (over far-peripheral V1) to regions associated with the FPN, including the mid orbitofrontal and inferior parietal regions of the FPN, as well as lateral portions of the DMN, and the insular portion of the CON. In contrast, far-peripheral representing V1 was preferentially connected (over central V1) to medial portions of the DMN (Figure 5).

To follow up on functional connectivity findings (Figure 4), we performed pairwise comparisons of cortical track terminations between the FPN to the central eccentricity segment of V1. Results indicate that, like our functional connectivity findings, there are also preferential structural connections between central V1 and the FPN compared to far-peripheral V1 (Figure 6).

### 3.3 Comparison of Functional and Structural Connectivity Patterns

To compare the patterns of functional and structural connections, we calculated a Dice Coefficient comparing a map of structural connectivity differences between central vs. far-peripheral connectivity where central V1 connections were stronger than peripheral connections to the analogous map of functional differences in the same set of vertices. Vertices included in this analysis included all vertices within FPN, CON, and DMN combined, as these were the vertices with information about structural connections to V1 in our analysis. The Dice Coefficient between structural and functional connectivity patterns was .330.

We then examined each of the three networks, individually for how similar the structural and functional connectivity findings were within the vertices of each functional network. Within the FPN, the Dice Coefficient (averaged across left and right hemisphere) between structural and functional connectivity patterns was .392. Within the CON, the Dice Coefficient (averaged across left and right hemisphere) between structural and functional connectivity patterns was .398. Within the DMN, the Dice Coefficient (averaged across left and right hemisphere) between structural and functional connectivity patterns was .444. These relationships indicate that the overall pattern of vertices with stronger connections to central V1 than peripheral V1 c is moderately consistent across modalities, a result consistent with prior work (Honey et al., 2009).

### 3.4 Correspondence between Functional Networks and Multi-modal Connectivity Patterns

Figure 7 shows regions where the central V1 segment had higher connectivity than the far-peripheral V1 segment in both the structural and functional comparisons. These regions include lateral frontal portions of the FPN, and small portions of other networks show overlap between the modalities (including the IFG portions of the DMN as defined from the Yeo et al., 2011 atlas in Figure 3).

We calculated the Dice coefficient to quantify the overlap between multi-modal regions of central dominance (as shown in Figure 7) and functional networks (FPN, DMN, CON). The Dice coefficient between the FPN and the multi-modal overlap was .165, DMN was .008, and CON was .004. This showed a relatively stronger overlap of the central V1 dominant network with the FPN than other networks.

## 4. DISCUSSION

Our goal was to better understand the brain network basis for interactions between sensory and higher-order information, especially how this differs between central vs. peripheral vision. Understanding the structural and functional underpinnings of these interactions is essential for understanding the processing differences between central and peripheral vision and for future work examining the plasticity of these systems.

Our approach compared structural and functional connections among different retinotopic eccentricities within V1 and large-scale functional networks (Fronto-Parietal Network (FPN), Cingulo-Opercular Network (CON), Default Mode Network (DMN)). Our results indicated that different visual eccentricities have different connectivity patterns to the rest of the brain, consistent with our previous data (Griffis et al., 2017) and data from other analyses (Buckner & Yeo, 2014). The present functional connectivity analyses replicated and extended previous findings on patterns of preferential connections between the central V1 eccentricity segment and the FPN (Griffis et al., 2017). Our structural connectivity analyses furthered the field’s understanding of the relationship between V1 and functional networks by describing the retinotopic pattern of structural connections. A comparison between structure and function showed moderate agreement, indicating that the functional connections are likely mediated by some direct structural connections. Further, the overlap of structural and functional findings (Figure 7) indicated the lateral frontal portions of the FPN and other nearby regions also responsible for attention and control (Inferior frontal gyrus (IFG)) made up more of the overlapping regions than the CON or DMN.

The present study found differences between the connection patterns of central and peripheral representations in V1. Since central and peripheral representations are still part of the same V1 cortical area, we would expect similarity in their connectivity patterns. Our results indicate that eccentricity differences in connection strength exist and are consistent with previously reported differences in information processing central and peripheral visual information. Central vision appears to be under more substantial top-down attentional control than peripheral vision (Chen & Treisman, 2008; Lu et al., 2002; Zhaoping, 2017). For example, stimuli presented within the peripheral visual field are more challenging to ignore than stimuli presented within the central visual field (Chen & Treisman, 2008). The current work suggests that this distinction may come from anatomical relationships to attentional networks.

### 4.1 Functional Connectivity

Our findings are consistent with prior work, specifically, preferential connections between central representing segments of V1 and regions belonging to the FPN (Griffis et al., 2017). These results suggest that frontal areas influence cognitive control mechanisms and primary visual processing areas, specifically central V1. On the other hand, the peripheral regions seemed to be preferentially connected more broadly across the cortex, with the specific exclusion of the FPN regions. The data provide further evidence to support the hypothesis of eccentricity dependent preferential connectivity of V1 to higher-order brain networks. One contribution in describing this connectivity is to extend previous work by Griffis et al., 2017, into a much larger dataset that was collected without fixation at rest, thereby improving the generalizability of the findings.

### 4.2 Structural Connectivity

Finding preferential structural connections between frontal regions to central V1, is consistent with our functional connectivity findings. These results support our hypothesis, based on functional connectivity findings, that the connections between V1 and brain regions associated with attentional control depend on eccentricity. As previously discussed, Markov and colleagues (2012) investigated direct structural connections in the macaque brain and found weak long-range connections between V1 and regions that may correspond to the human FPN. These connections were projections *from* V1 to frontal and parietal regions (areas F5, 81, and 7A), rather than projecting *to* V1. These results could indicate that the structural connections observed here are bottom-up connections that provide visual information to direct cognitive control within the FPN. However, previous work in macaques has not found major white matter tracts connecting the occipital lobe to the frontal lobe (Takemura et al., 2016). Further, the attention system of the macaque is different from that in humans (Patel et al., 2015). Thus, the prior macaque literature provides limited insight into the structural connections between V1 and FPN in humans, so the direction of these connections is still unclear. Although diffusion tractography methods can conflate crossing fibers, tractography showed strikingly similar effects to our functional connectivity results, bolstering the plausibility of an interpretation of a direct connection between lateral frontal regions to V1. Describing this connectivity extends the knowledge of functional connectivity between V1 and functional networks and improves our understanding of the structural underpinnings of these functional connections.

### 4.3 Overlap Between Structural and Functional Connections and their Relationship to Functional Networks

Structural connectivity patterns and functional connectivity patterns showed correspondence. Because diffusion tractography is typically interpreted as a direct structural connection, the functional connections between regions that also showed a structural connection likely reflect (at least in part) direct structural connections. These regions that showed centrally-weighted structural and functional connections are shown in Figure 7, and include inferior frontal gyrus.

A direct, long-range connection between IFG and central vision representations in V1 may be related to the importance of speed in attentional control. For example, visual information needs to be processed quickly to impact attention selection. Thus, attentional control from the FPN to central vision representations in V1 would be improved by direct structural connections. Processing of central vision information in FPN circuits would also be improved by direct structural connections. A large body of work supports top-down and bottom-up effects on visual processing (Gandhi et al., 1999; Somers et al., 1999; Tootell et al., 1998; Yeshurun & Carrasco, 1998; Zhaoping, 2017). The present study contributes to this field by demonstrating the eccentricity dependent nature of the relationship between V1 and higher-order brain regions.

### 4.4 Relationship between central vision processing and attention

Complex biological systems are often driven by separate control mechanisms with distinct functional properties (Dosenbach, Fair, Cohen, Schlaggar, & Petersen, 2008). During cognitive operations, information processing appears to rely upon the dynamic interaction of brain areas as large-scale neural networks including FPN, CON, and DMN. FPN supports executive functions by initiating and adjusting top-down control (Dosenbach et al., 2008). The CON supports salience-related functions and provides stable control over entire task epochs. The suppression of DMN is critical for goal-directed cognitive processes (Spreng, Stevens, Chamberlain, Gilmore, & Schacter, 2010). Cooperation among these top-down control systems of the brain is necessary for controlling attention, working memory, decision making, and other high-level cognitive operations (Hellyer et al., 2014; Keller et al., 2015; Raichle, 2015; Ray et al., 2019; Spreng, 2012; Vincent, Kahn, Snyder, Raichle, & Buckner, 2008).

FPN includes regions such as the intraparietal sulcus that play an essential role in goal-directed cognitive functions (Spreng et al., 2010), and both spatial and non-spatial visual attention (Giesbrecht, Woldorff, Song, & Mangun, 2003; Scolari, Seidl-Rathkopf, & Kastner, 2015). The role of central vision in visual processing and object recognition and the need to inhibit distractors in the visual field could (speculatively) be an evolutionary reason central vision representations are preferentially connected to attention regions, including the FPN. Contributions from high-order cognitive areas, like the FPN, help the brain decide which visual areas will be prioritized for visual attention (Scolari et al., 2015).

### 4.5 Limitations and Future Directions

Our study has several methodological limitations that we will discuss here. Our study used only healthy young adults from the HCP dataset, which could influence our findings’ generalizability. Future work should include individuals from across the lifespan.

The task that participants completed during the HCP protocol were quite distinct from the previous dataset’s task (Griffis et al., 2017). Here, participants rested quietly, and though their instruction was to keep eyes open, no one assessed if the participants’ eyes were open or closed. In contrast, the Griffis dataset included data from the rest period between blocks of a task, and eye position and lid opening were confirmed via eye-tracking. The fact that these data closely follow each other extends the possible interpretations of the original dataset: the distinction between peripheral and central V1 connectivity generalizes to a new task context.

It should also be acknowledged that functional connectivity can be influenced by attention (Gratton et al., 2018; Griffis et al., 2015; Salehi et al., 2020). In both the work by Griffis and colleagues (2017) and the current study’s resting-state scan, a fixation cross presented on a screen at the end of the bore, and participants were scanned while inside the MRI bore. Participants may, therefore, have been allocating more attention toward the visual space in the center (the screen) than the periphery (the bore). However, the fact that we observed complementary effects in the structural data indicates that these data are likely not due to transient states of attention and are likely to represent biological organization.

We acknowledge that large veins near posterior occipital cortex area V1 could impact our functional connectivity measurements in this area (Winawer, Horiguchi, Sayres, Amano, & Wandell, 2010). However, we performed extensive pre-processing to reduce the impact of vessels on the results. In addition, the voxel size of our resting state scan is small (2mm isotropic); this higher resolution should mitigate contributions from nearby veins due to partial voluming effects (Schira, Tyler, Breakspear, & Spehar, 2009).

Functional connectivity strengths between V1 to the lateral frontal cortex are on the order of r=0.1 (Figure 4). While very reliable, these magnitudes are not as large as connections to other areas, for example, portions of the occipital lobe. Functional connectivity magnitudes are always influenced by the preprocessing done to obtain them. In this case, we regressed out the mean signal and regressed out white matter and CSF. While this practice decreases the mean correlation strength (Shirer, Jiang, Price, Ng, & Greicius, 2015; Weissenbacher et al., 2009), it also improves across-subject reliability (Burgess et al., 2016). The debate about this practice, now a decade long, has focused on the interpretability of negative correlations, which we do not do here; our inferences are based on differences in correlations across brain areas.

DWI-based tractography produces similar results to tracer methods (Donahue et al., 2016); however, probabilistic tractography indirectly traces axon bundles by modeling the path of most restricted water movement and then estimating white matter tracts. Fibers that cross, fan, or converge pose problems for accurately estimating white matter tracts (Johansen-Berg & Rushworth, 2009). One way to improve track estimation is by modeling multiple angular compartments (e.g., ball-and-stick model) and using greater than 30 diffusion directions (i.e., 95, 96, and 97 directions in the present study), both of which were used in the present study (Behrens et al., 2007). Connections described in tractography are non-directional in that no determination of the direction of signaling is acquired. Therefore, the current study cannot interpret the direction (top-down versus bottom-up processing) of the described connections outside of the context of prior tracer studies.

Although the present tractography and functional connectivity analyses aim to measure connections between eccentricity segments of V1 and functional networks, they are inherently different modalities, including but not limited to differences in the measurement of direct and indirect connections between regions. Structural tractography analysis only identifies direct connections, and functional connectivity analysis can identify direct and indirect connections. Therefore, the comparison between them is limited in scope. Since tractography describes direct connections between brain regions, inconsistencies, where functional connectivity is present but structural connectivity is not, could be due to indirect connections (Honey et al., 2009). However, measuring both structural and functional connectivity provides valuable information to understand the relationship between brain regions that cannot be derived from one modality alone.

Future work could help determine the direction of the observed connections and further describe the complexity of direct and indirect connections between V1 and functional networks, and could also examine extrastriate regions that are retinotopically mapped. While we only studied participants with healthy vision, future work should include participants with low vision to investigate possible connectivity changes related to vision loss. This work could serve as a baseline for these low vision studies. Future research could also help inform the plasticity of the described connections in the context of visual training and vision loss.

### 4.6 Conclusions

There is a reliable preference for central representations in V1 to be more strongly connected to the frontoparietal network than peripheral visual representations. This is particularly true for the lateral frontal portions of that network and the Inferior Frontal Gyrus, and is true for structural as well as functional connections. This implies that the functional relationship between central V1 and frontal regions is built, at least in part, upon direct, long-distance connections. Understanding how V1 is functionally and structurally connected to higher-order brain areas contributes to our understanding of how the human brain processes visual information and forms a baseline for understanding any modifications in processing that might occur with training or experience.

In summary, the work’s main contribution is a greater understanding of higher-order functional networks’ connectivity to the primary visual cortex (V1). Centrally-representing portions of V1 are connected to some frontal cortical regions, including portions of the FPN, both functionally and structurally. The lateral frontal regions where connection differences overlap between structural and functional data have been associated with attention in previous work. This suggests that the central representations of V1 are more tightly coupled to some brain regions involved in attention and cognitive control. Understanding how V1 is functionally and structurally connected to higher-order brain areas contributes to our understanding of how the human brain processes visual information and forms a baseline for understanding any modifications in processing that might occur with training or experience.

## Acknowledgments

Thanks to Jen Robinson for technical help with tractography analysis, Utkarsh Pandey for data cleaning, Joe Griffis, Wes Burge, and Rodolphe Nenert for code for functional connectivity and regions of interest, John Paul Robinson and Ravi Tripathi for assistance with performing high performance computing, and the members of the Visscher Lab. Thanks to funding from NIH U01 EY025858 and NIH/NINDS T32NS061788-12 07/2008 – 0. Data were provided by the Human Connectome Project, WU-Minn Consortium (Principal Investigators: David Van Essen and Kamil Ugurbil; 1U54MH091657) funded by the 16 NIH Institutes and Centers that support the NIH Blueprint for Neuroscience Research; and by the McDonnell Center for Systems Neuroscience at Washington University.

## Data and code availability statement

Data is available through the Human Connectome Project (900 subject release):

http://www.humanconnectomeproject.org/data/

Code for preprocessing and analysis has been made available at: https://github.com/Visscher-Lab/V1_eccentricty_HCP_analysis/

## Supplemental figure

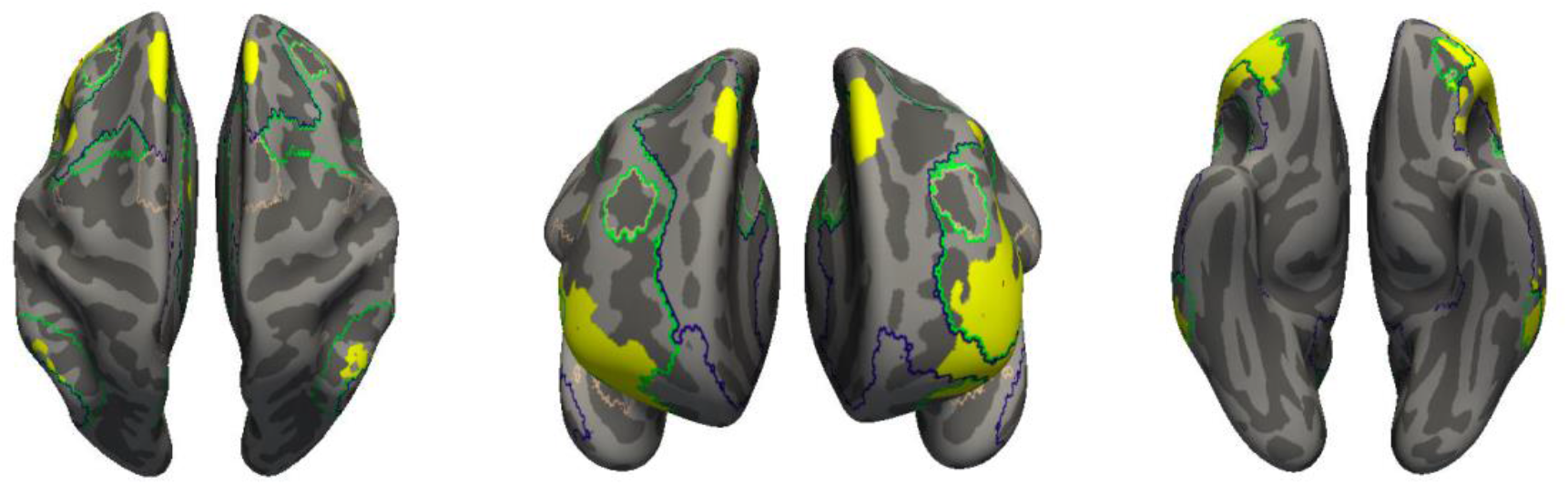
Additional views of Figure 7.

## Notes

### Competing Interest Statement

The authors have declared no competing interest.

### Summary of Updates

Revisions based on comments from eLife reviewers.

